# Extensive aquatic subsidies lead to territorial breakdown and high density of an apex predator

**DOI:** 10.1101/2021.03.29.437596

**Authors:** Charlotte E. Eriksson, Daniel L.Z. Kantek, Selma S. Miyazaki, Ronaldo G. Morato, Manoel dos Santos-Filho, Joel S. Ruprecht, Carlos A. Peres, Taal Levi

## Abstract

Energetic subsidies between terrestrial and aquatic ecosystems can strongly influence food webs and population dynamics. Our objective was to study how aquatic subsidies affected jaguar (*Panthera onca*) diet, sociality, and population density in a seasonally flooded protected area in the Brazilian Pantanal. The diet (n = 138 scats) was dominated by fish (46%) and aquatic reptiles (55%), representing the first jaguar population known to feed extensively on fish and to minimally consume mammals (11%). These aquatic subsidies supported the highest jaguar population density estimate to date (12.4 per 100 km^2^) derived from camera traps (8,065 trap nights) and GPS collars (n = 13). Contrary to their mostly solitary behavior elsewhere, we documented social interactions previously unobserved between same-sex adults including cooperative fishing, co-traveling, and play. Our research demonstrates that aquatic subsidies seen in omnivores can be highly influential to obligate carnivores leading to high population density and altered social structure.

## Introduction

Energetic subsidies through the transfer of resources between terrestrial and aquatic ecosystems can strongly influence food webs and population dynamics (Polis, Anderson and Holt, 1997). The transfer of energy from large emergences of aquatic insects to terrestrial systems is the canonical example of this phenomenon (Nakano and Murakami, 2001), but fish-consuming terrestrial vertebrates can also play an important role. Salmon systems famously support hyperabundant bear (*Ursus arctos*) populations with modified social dynamics due to the need to tolerate conspecifics at point sources (e.g., waterfalls and streams) where salmon become accessible (Egbert and Stokes, 1974). Similarly, Rose and Polis (2016) found coastal coyote (*Canis latrans*) populations subsidized by marine resources to be on average 4-5 times more abundant than inland populations. Several other omnivorous terrestrial carnivores forage in marine or freshwater systems such as gray wolves (*Canis lupus*), raccoons (*Procyon lotor*) and arctic foxes (*Vulpes lagopus*) (Carlton and Hodder, 2003), but aquatic subsidies in obligate terrestrial carnivores are thought to be rare. Of the obligate terrestrial carnivores, jaguars (*Panthera onca*) have been most frequently linked to aquatic resources such as caiman and chelonians (da Silveira *et al.*, 2010), although terrestrial mammals still dominate their diet.

A resource-rich environment may enable multiple individuals to share a common resource, which typically results in higher animal densities, smaller home-ranges and increased social interactions among individuals according to the resource dispersion hypothesis (Macdonald, 1983). Most large felids, including the jaguar, are considered solitary species with social interactions generally limited to courtship and territorial disputes (Sunquist and Sunquist 2002). However, the degree of intraspecific tolerance and sociality can change with resource availability (Elbroch & Quigley, 2017). The Pantanal wetlands in central South America harbors abundant terrestrial and aquatic jaguar prey resources. Here, jaguars reach high population densities (Soisalo and Cavalcanti, 2006) and use smaller home-ranges than elsewhere in their range (Morato et al., 2016), but spatial tolerance of conspecifics, defined by overlapping home-ranges, seem to vary. Azevedo and Murray (2007) found that jaguars maintained exclusive territories while Cavalcanti and Gese (2009) found extensive overlap among males but not among females. Consumption of aquatic prey, mainly caiman (*Caiman yacare*), varies among study areas from 20-30% with terrestrial mammals still constituting the majority of the diet in all prior research (Fig. 2C). In addition, domestic cattle are abundant and make up a considerable portion of the jaguar diet in the southern Pantanal where most research has been conducted to date. Thus, it is currently unclear whether aquatic subsidies have altered the abundance or social structure of these jaguar populations, as has been demonstrated with other omnivorous carnivores.

**Figure 1.**
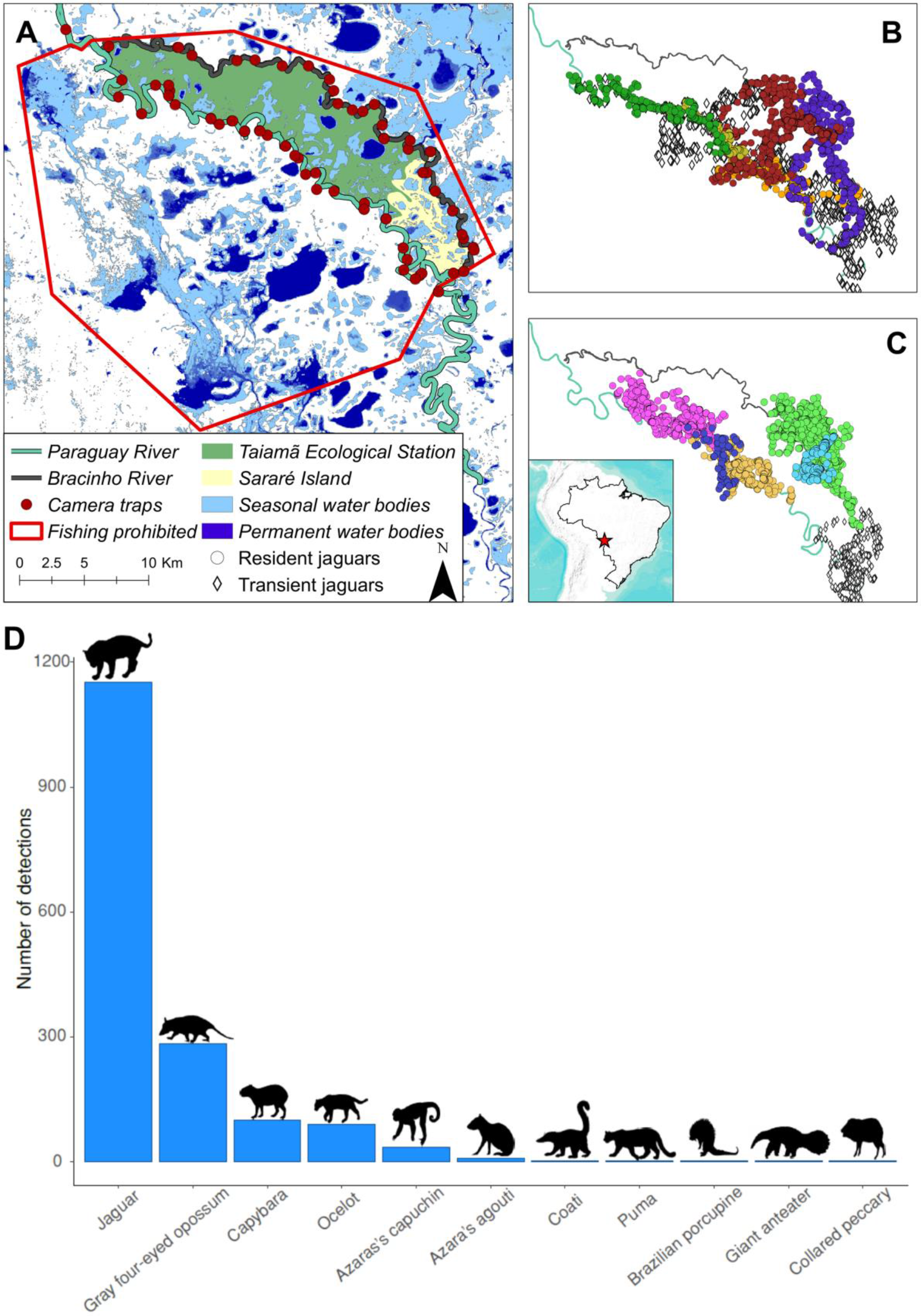
(A) Map of study area at Taiamã Ecological Station within the Pantanal wetlands in Mato Grosso, Brazil. (B) GPS locations of six resident males and one transient. C) GPS locations of five resident females and one transient. D) Number of detections of medium and large mammal species at Taiamã Ecological Station using camera traps from 2014-2018.

**Figure 2.**
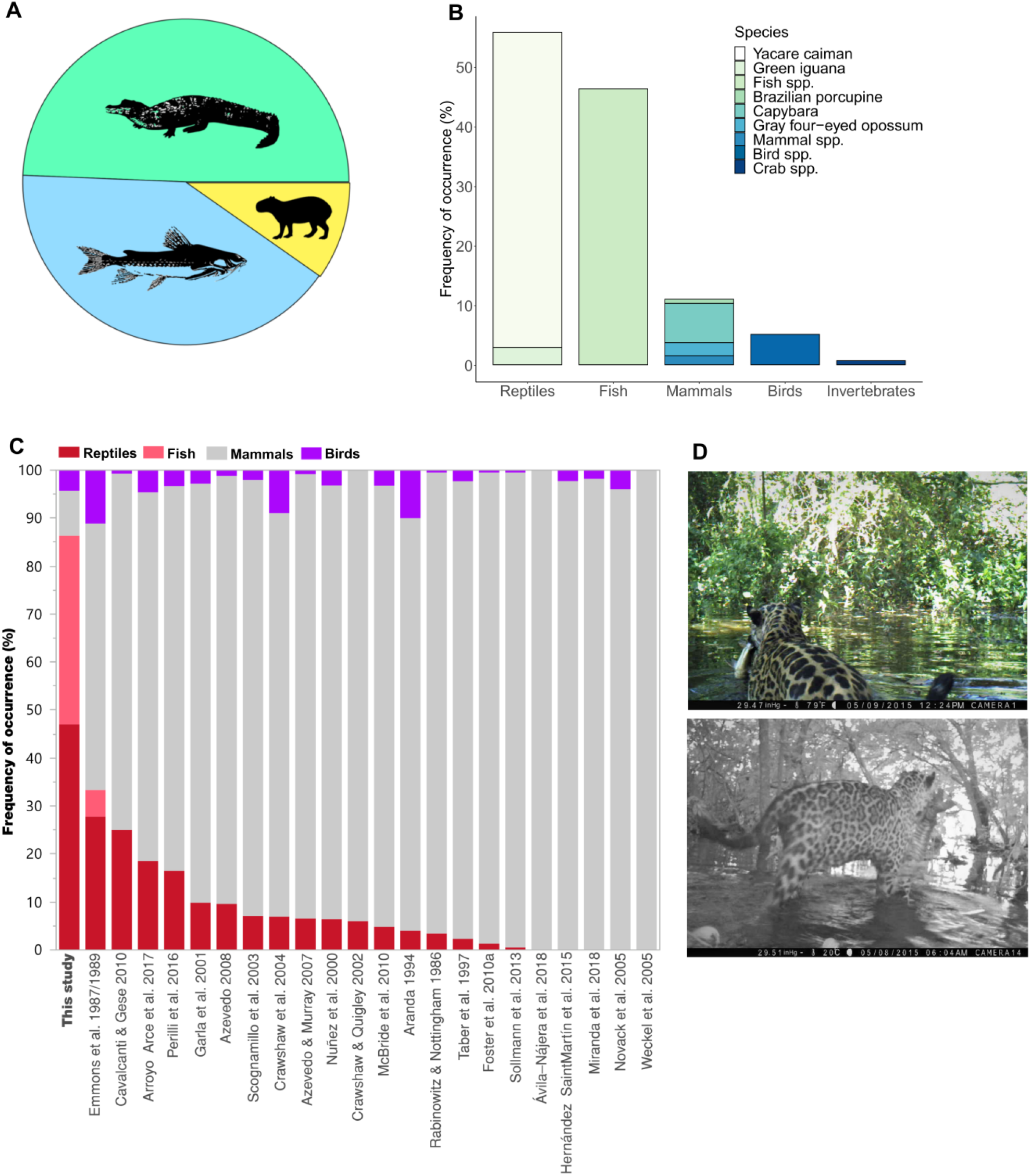
Frequency of occurrence of prey found in jaguar scats (n = 138) in Taiamã Ecological Station between 2013 and 2018. (A) Main food groups: reptiles, fish, and mammals. (B) All identified prey items. (C) Frequency of occurrence of vertebrate prey groups found in diets throughout the geographic range of jaguars. Only peer-reviewed studies with a sample size of at least 20 scats or kill sites were included (See Appendix S4: Table S1 for more information). (D) F8 carrying a tiger catfish (*Pseudoplatystoma fasciatum*) and M4 after catching a thorny catfish (*Doradidae* spp.).

In contrast to the southern Pantanal, the ecology of jaguars in the wetter northern portion (i.e., within the Brazilian state of Mato Grosso) remains largely unstudied. Taiamã Ecological Station (hereafter, Taiamã), a small and seasonally flooded protected area (115.55 km^2^), was suggested to have an unusually high jaguar density based on frequent jaguar sightings (Kantek and Onuma, 2013). A large presence of jaguars in this area is further made obvious by abundant jaguar footprints, scats, scratch marks on trees and large caiman carcasses scattered along the river margins. Taiamã (16°50’ S and 57°35’ W, Fig. 1A) was established in 1981 to protect this unique wetland ecosystem and its high diversity and abundance of fish species (ICMBIO 2017). Importantly, there are no human settlements or cattle ranching within or in the immediate vicinities of Taiamã and fishing is strictly prohibited (ICMBIO 2017). Our objectives were to study the ecology of jaguars at Taiamã to determine if the presence of abundant aquatic resources affect their diet, social behavior and ultimately population density, and to place these findings in the context of previous literature on jaguar ecology. To accomplish these objectives, we collected jaguar scats for dietary analysis, estimated jaguar density and documented social interactions using camera traps and telemetry data from 13 jaguars. This research details a rare example of how allochthonous resource subsidies can be so prolific that they influence the behavior and social organization of an apex predator.

## Methods

### Study Area

The study area encompassed Taiamã Island and Sararé Island bounded by the Paraguay and Bracinho rivers within the Brazilian Pantanal (Fig. 1A). The area is characterized by permanent lakes and seasonally flooded grasslands and riparian gallery forests. The climate is defined by a dry (April-September) and wet season (October-March) with an average annual precipitation of 1,200-1,300 mm. The maximum river level occurs between February-June and the lowest in August-November. The fluctuating water levels substantially alter the availability of terrestrial vs aquatic habitats (Fig. 1A) and determine local ecological patterns and processes, including the distribution and abundance of wildlife (Mamede and Alho, 2006).

### Camera dataset

We carried out three camera trapping surveys between 2014 and 2018. The absence of roads or trails coupled with the prolonged flood pulse and the high jaguar density presented unique logistical challenges for field work and prevented any movement by car or foot. All camera deployments were therefore boat-based and restricted to the major waterways and, thus, prevented the use of a systematic camera grid. We used unbaited Bushnell (Trophy Cam 119636) and Browning (Dark Ops BTC-6) camera traps separated by approximately 2 km along the Paraguay and Bracinho rivers. The two waterways were parallel to each other and separated by 4-6 km (Fig. 1A). Camera availability varied across the years. In 2014 and 2015 39 and 27 single cameras were set to video mode (20 sec/video) and operated for 56 (August-October) and 252 days (March-November), respectively. In 2018 one camera was set to video and the other paired camera was set to take photos (three per burst) for 65 days (August-October) at 20 camera stations. Cameras were set approximately 50 cm off the ground on trees and checked every three weeks. The study area encompassed an area of approximately 236.7 km^2^, which is larger than a jaguar home range in this area (Morato et al., 2016) as recommended for accurate density estimation (Tobler and Powell, 2013). Individual jaguars were identified in each photo or video based on their unique coat patterns. The camera detections were collapsed into weekly occasions for density estimation. The 252-day long camera deployment in 2015 was split into two sessions to assume demographic closure resulting in four sessions of 8, 18, 18 and 10 weekly occasions each. Camera trap sampling details are summarized in Appendix S1: Table S1.

### Telemetry dataset

Seven male and six female jaguars were captured between December 2011 and April 2016 as part of concurrent studies (Morato *et al.*, 2016) (SISBIO #30896-3). Jaguars were captured using foot snares and immobilized with tiletamine and zolazepam (10 mg/kg). Two of the jaguars were fitted with VHF radio-collars and the remaining (n = 11) were fitted with Lotek-Iridium GPS collars programmed to record location data every hour and to automatically drop-off the animal after 400 days.

### Density estimation

A fundamental issue with density estimation is how to deal with lack of geographic closure given that some fraction of detected individuals primarily lives outside the study area. Spatially explicit Capture-Recapture models (SCR; Borchers and Efford 2008; Royle and Young 2008) have become a preferred method for density estimation over conventional non-spatial capture-recapture techniques that are sensitive to the arbitrarily defined effective sampling area. While SCR models present a useful advance over conventional capture-recapture models, they assume isotropic (i.e., circular) home ranges such that the detection probability of each individual falls off equally with distance from its activity center. Our telemetry data revealed that the direction of jaguar movement at Taiamã follows the rivers closely and the home ranges are therefore elongated and highly anisotropic (Fig. 1B and C). SCR models may still perform well if sufficient detectors are placed in two dimensions capturing both the major axis of movement (along the rivers in our case) and the minor axis (perpendicular to the rivers) (Efford, 2019). The bias can also be corrected by applying anisotropic detection models if all individual home ranges are directionally aligned (Murphy *et al.*, 2016). Since our camera deployment was boat-based and restricted to the rivers, we were unable to effectively capture jaguar movement in the minor axis, and telemetered jaguars did not consistently share the same home range orientation (Fig. 1B, C). Applying a SCR model to this unusual dataset where the detectors are aligned with elongated home ranges would therefore lead to an overestimation of the spatial scale parameter and consequently underestimate density (Efford, 2019).

For data-rich applications where a representative portion of the population has been telemetered, the extent of geographic closure violations can be corrected for directly (Ivan, White and Shenk, 2013) to estimate population density without the restrictive assumptions of circular home ranges or the size of the effective sampling area. This method uses a conventional Huggins closed-capture model augmented by telemetry data to estimate the proportion of time telemetered individuals spend within the sampling area (i.e., residency) to directly correct for closure violations. We used this method with the modification of allowing capture probability to vary by sex and included a random effect on residency by individual jaguar so that one individual’s data would not dominate the estimate (full methods and JAGS code in Appendix S2).

### Diet analysis

Jaguar scats were collected opportunistically during field work. Undigested remains found in the scats were isolated by rinsing in a fine mesh, dried, and identified to the lowest taxon possible using a dissecting scope. We compared our frequency of occurrence diet estimates to other published work that used either scat or kill site analysis. For comparative purposes, we grouped prey items into ‘mammals’, ‘reptiles’, ‘fish’ and ‘birds’ and standardized frequency of occurrence values to sum to 100%.

### Social interactions among jaguars

We investigated shared space use among the resident jaguars collared in 2014-2015 (n=9), by quantifying the proportion of overlap at the 95% individual utilization distribution using the Autocorrelated Kernel Density Estimator (AKDE) in the R package ‘CTMM’ (Calabrese, Fleming and Gurarie, 2016). Because jaguars exhibiting home-range overlap can still be avoiding each other temporally, we analyzed the frequency of social interactions by assessing simultaneous locations from both GPS data and camera data. GPS interaction rates were calculated using the package wildlifeDI (Long and Nelson, 2013) in R and defined as the number of times two GPS-collared jaguars were within 200 m of each other (Cavalcanti and Gese, 2009; Elbroch and Quigley, 2017). We defined camera interaction rates as the number of times two jaguars were observed together or within 30 min of each other. We collapsed interactions between individuals per day such that multiple contacts within the same day were not counted as new interactions for both the GPS and camera data.

## Results

We operated 59 camera stations for 8,065 days between 2014 and 2018 (Appendix S1: Table S1; Fig. 1A). Jaguars were detected at 95% of the cameras (n = 56) and were the most frequently detected mammal (Fig. 1D). We obtained 1,594 videos of jaguars (excluding the photos from the paired cameras in 2018), representing 385 individual weekly detections with the number of detections per individual ranging from one to 34 (mean 5.6 ± 5.9 SD). We detected 12 out of the 13 telemetered jaguars with our camera traps. We detected an additional 29 females, 21 males, two individuals of unknown sex, and four cubs, representing a total of 69 unique individuals. The maximum number of unique jaguars captured by one camera was 15 across all 3 years including 9 just in 2015. Both right and left flanks were obtained for most of the individuals using both our video setup in 2014 and 2015 and paired camera setup in 2018. One jaguar was captured only on her left side but was included in the dataset since the closest female with only a right-side capture was detected 20 km away and had a distinct rosette pattern. We were therefore able to retain all identified jaguars in the analyses. The highest minimum number of jaguars known to be present within the study area within one camera session was 40 (Appendix S1: Table S1).

The telemetered jaguars spent on average 96% of the time within the study area (Fig. 1B, C) Estimated jaguar density was 12.4 per 100 km^2^ and mean capture probability per weekly occasion was 0.18 (0.15 - 0.21 Bayesian Credible Interval; BCI) for females and 0.27 (0.24 - 0.31 BCI) for males. Density was similar across the three dry season sessions at 12.3 (10.1 - 14.4), 11.9 (10.1 - 12.9) and 11.3 (9.4 - 13.1) jaguars per 100 km^2^, in session 1, 3, and 4, respectively. The wet season estimate from session 2 was 14.3 (12.2 - 15.4) jaguars per 100 km^2^ (Appendix S1: Table S2), even though some 40% of the Taiamã land area becomes seasonally flooded.

### Jaguar diet

We identified nine prey items in 138 jaguar scats collected from 2013-2018. The jaguar diet was dominated by three taxonomic groups: reptiles (55%), fish (46%) and mammals (11%) (Fig. 2A). These findings were corroborated by camera data showing jaguars feeding on fish and caimans, and other field observations such as large caiman skulls and fish carcasses scattered along the rivers. Fish remains found in the scats could not be identified to species, however, the camera data revealed jaguars capturing thorny catfish (*Doradidae*) (Fig. 2D), pacu (*Piaractus mesopotamicus*), red-bellied piranha (*Pygocentrus nattereri*), and large tiger catfish (*Pseudoplatystoma fasciatum*) (Fig. 2D; see Appendix S2: Fig.S1 for jaguar fishing behavior). Most mammals consumed were semi-aquatic capybaras (*Hydrochoerus hydrochaeris*). Rare prey items included green iguana (*Iguana iguana*), Brazilian porcupine (*Coendou prehensilis*), freshwater crab, and gray four-eyed opossum (*Philander opossum*) (Fig. 2B).

### Social interactions

All resident GPS-collared males had overlapping 95% AKDE home ranges, except for M3 and M4 (Appendix S2: Fig.S2A). M1 overlapped from 17.6 to 25.5% with M2 and M3, respectively, while M2 and M3’s home ranges overlapped by 64.7%. M5 overlapped with M1 by 7.6% and 16.1% with M3. The 95% AKDE home range also overlapped extensively between GPS-collared females (Appendix S2: Fig.S2B). F2’s home range overlapped by 58.6% with F1 and the home ranges of F3 and F4 overlapped to 21.6%. F1 and F2 were unrelated while F3 and F4 were mother and daughter (Kantek et al., 2021). Including jaguars detected only by camera, between 4-15 individuals were detected within the home ranges of telemetered jaguars indicating even higher spatial tolerance than observed with GPS data alone.

We documented 80 independent social interactions between adult jaguars based on the camera (n=40) and GPS data (n=40). The majority of the interactions were between males and females (M-F 85%), and involved 29 individual jaguars (14 males and 15 females). We documented 12 interactions of the same sex (one F-F and 11 M-M) consisting of 10 individuals (two females and eight males). Two GPS-collared males, M2 and M1, moved continuously within approximately 41-182 m of each other for two days. Eight months later they were captured with the same camera 5 min apart. M2 moved continuously with another GPS-collared male (M3) for 10 hours within 4-38 m of each other, again two days later (2 hours; 14 m apart) and were close again two weeks later (1 hour; 159 m). None of the three males were related (Kantek et al., 2021). Two other males (M7 and M21) were seen fishing together (Fig. 3D) and walking past another camera together on two separate occasions. Two males (M29 and M27) spent 30 min in front of a camera “playing” (Fig. 3E and F). Two females (F2 and F4) were documented in the same location 12 min apart (Fig. 3G). We lack information on potential relatedness among the interacting uncollared jaguars.

**Figure 3.**
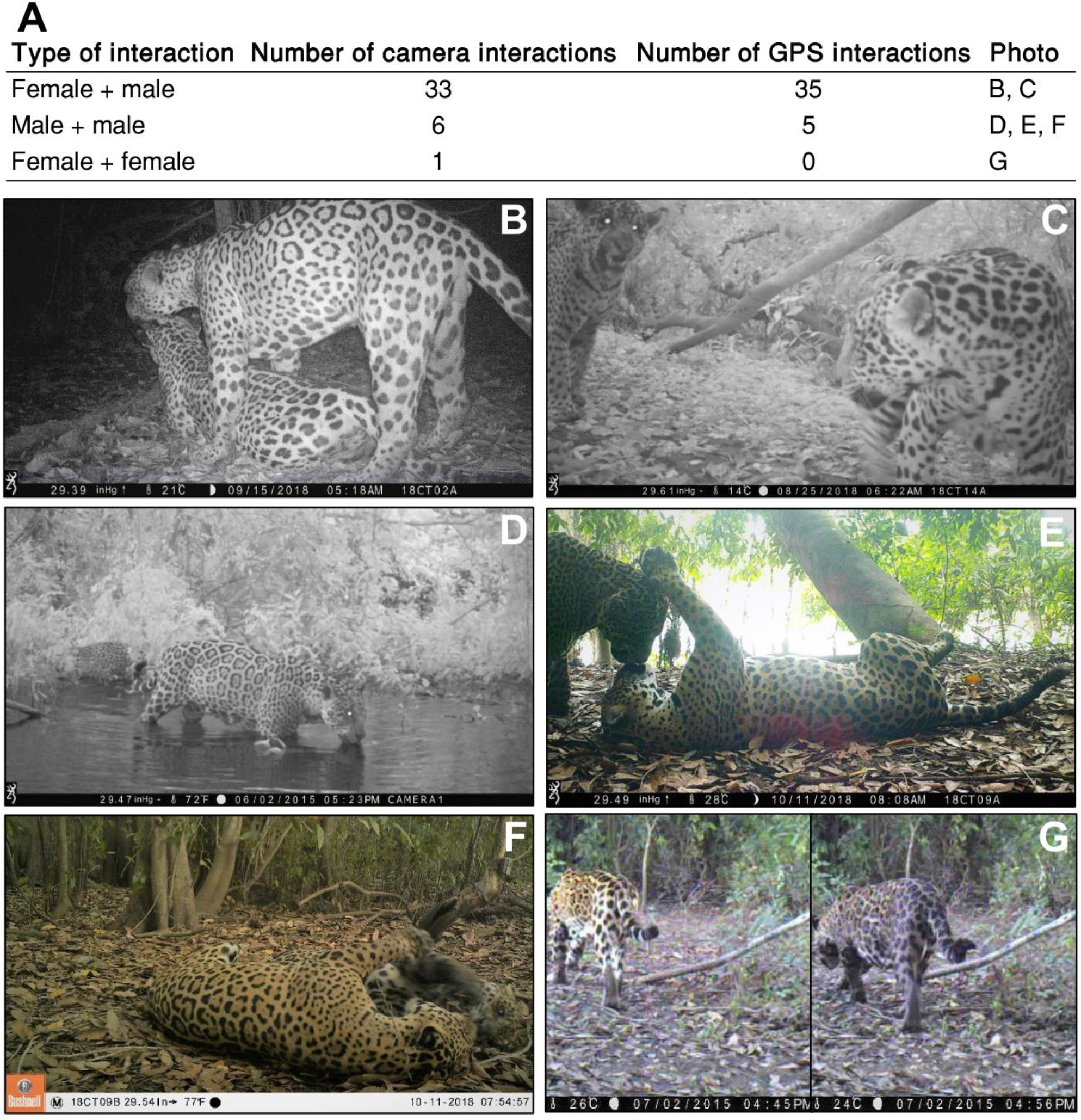
(A) Number and type of social interactions among adult jaguars determined by GPS data and camera trap data. (B-C) Male and female mating behavior, (D) Two adult males fishing together, (E-F) two adult males playing, (G) Two females captured with the same camera 12 minutes apart.

## Discussion

We have reported on an extraordinarily high density for a large-bodied apex predator that is extensively sustained by aquatic subsidies. Jaguars in Taiamã have by far the most aquatic diet and the least mammal consumption of any previously studied population. Although jaguars in Taiamã consume more aquatic reptiles than has ever been observed, it is fish consumption that makes this population truly unique (Fig. 2C). As far as we know, this is the most piscivorous diet of any large felid and among the most for any terrestrial hypercarnivore. Even the famous tigers (*Panthera tigris*) in the Sundarbans mangroves of India still consume mostly terrestrial mammals (Aziz, Islam and Groombridge, 2020). The small fishing cat (*Prionailurus viverrinus*) may be the most similar example of a piscivorous felid akin to the jaguars at Taiamã (Ganguly and Adhya, 2020). Although jaguars are commonly associated with rivers (Morato *et al.*, 2018) and local knowledge indicates that jaguars prey on fish, the only published records of fish consumption include anecdotal observations of jaguars fishing summarized in Gudger (1946) and more recently Emmons (1987) found fish in two jaguar scats. However, localized fish consumption may be more common than is currently appreciated. We are aware of one unpublished study from a coastal island in Northern Brazil that also indicates substantial reliance on fish (Vergara, 2011). Given limited previous research in this remote roadless region, it is possible that what we have discovered at Taiamã is broadly representative of jaguar foraging ecology within this highly flooded portion of the Pantanal.

These extensive aquatic subsidies support perhaps the highest density jaguar populations described to date (12.4 jaguars per 100 km^2^). Even just the 13 *telemetered* individuals that we knew were all present in 2015 with on average 96% of GPS locations contained within the study area (236.7 km^2^) would suggest a density of approximately 5.4 per 100 km^2^ without considering the additional 56 individuals detected with cameras. Just this density estimate from telemetered individuals is comparable or exceeds other high jaguar density estimates such as 4.5 jaguars/100 km^2^ from the Peruvian Amazon (Tobler et al., 2018), 4.4 jaguars/100 km^2^ in the Venezuelan Llanos (Jędrzejewski *et al.*, 2017), and 6.6 to 6.7 jaguars/100 km^2^ in the southern Pantanal (Soisalo and Cavalcanti, 2006).

According to the resource dispersion hypothesis, abundant resources can lead to a breakdown in territoriality and increased social tolerance among individuals (Macdonald, 1983). Aquatic subsidies are uniquely capable of providing the requisite levels of resource abundance because their ability to rapidly generate energetically inexpensive heterotherms in productive aquatic habitats. The association of aquatic subsidies with high population density and increased social tolerance of jaguars at Taiamã is similar to that observed with brown bears in systems with high salmon abundance. Evidence of high social tolerance at Taiamã includes highly overlapping home ranges, substantial time spent co-traveling, and numerous social interactions directly observed by video monitoring (Fig. 5). The most notable social interactions include social play and cooperative fishing behavior. The social play among adult animals observed here is rarely documented and is thought to occur when all biological needs have been met (Bekoff, 1972). The incident of play between M27 and M29 started with passive submissive patterns such as rolling over, facial rub, and face-paws (Fig. 5E, F). The play then transitioned to play-fighting with wrestling, soft bites, and mounting. Cooperative fishing occurred among a female and her older cub as well as two adult males that were also observed co-traveling on three occasions (Fig.5D; Appendix S2: Fig.S1). We cannot determine whether this form of cooperative fishing improved fish capture efficiency, or whether co-fishing jaguars simply tolerated the close proximity of each other without deriving any additional prey-acquisition benefits.

Our study demonstrates that aquatic subsidies, frequently described in omnivores, can be influential to the ecology and behavior of obligate carnivores as well. This work further sheds new light on the relationship between carnivore socioecology and prey abundance, showing that the modal pattern of solitary behavior observed in the majority of hyper-carnivores can be broken given the nature, productivity, and distribution of the resource base. Further research is needed to understand the role jaguars play in linking aquatic and terrestrial food webs. Potential implications of this highly-subsidized population for terrestrial prey are highly divergent including either the suppression of terrestrial prey via apparent competition (Holt, 1977) or release from predation pressure through aquatic diet specialization (i.e. apparent mutualism; (Holt and Bonsall, 2017). Finally, this research demonstrates the flexibility and context-dependence of animal ecology and behavior such that even well-studied charismatic vertebrates continue to surprise us.

## Acknowledgments

This study was funded by Helge Ax:son Johnson Foundation and Oregon State University. We thank ICMBio and UNEMAT for facilitating this research. We thank Manaav Kamath, Aurea Ignácio, Claumir Muniz, Amabile Furlan and Derrick Campos for logistical support and field work assistance, as well as Gerlane Costa for help with prey remain identification.

## Notes

### Competing Interest Statement

The authors have declared no competing interest.

## Literature cited

Alho, C. (2008) ‘Biodiversity of the Pantanal: response to seasonal flooding regime and to environmental degradation’, Brazilian Journal of Biology, 68(4), 957–966.

Azevedo, F. C. and Murray, D. L. (2007) ‘Spatial organization and food habits of jaguars (*Panthera onca*) in a floodplain forest’, Biological Conservation, 137(3), pp. 391–402.

Aziz, M. A., Islam, M. A. and Groombridge, J. (2020) ‘Spatial differences in prey preference by tigers across the Bangladesh Sundarbans reveal a need for customised strategies to protect prey populations’, Endangered Species Research. Inter-Research Science Center, 43, pp. 65–74.

Bekoff, M. (1972) ‘The development of social interaction, play, and metacommunication in mammals: An ethological perspective’, The Quarterly Review of Biology, 47(4), pp. 412–434.

Borchers, D. and Efford, M. (2008) ‘Spatially explicit maximum likelihood methods for capture– recapture studies’, Biometrics, (64), pp. 377–385.

Calabrese, J. M., Fleming, C. H. and Gurarie, E. (2016) ‘ctmm: an R package for analyzing animal relocation data as a continuous-time stochastic process’, Methods in Ecology and Evolution, 7, pp. 1124–1132.

Carlton, J. T. and Hodder, J. (2003) ‘Maritime mammals: terrestrial mammals as consumers in marine intertidal communities’, Marine Ecology Progress Series, 256, pp. 271–286.

Cavalcanti, S. M. and Gese, E. M. (2009) ‘Spatial ecology and social interactions of jaguars (*Panthera onca*) in the southern Pantanal, Brazil’, Journal of Mammalogy, 90(4), pp. 935–945.

Efford, M. (2019) ‘Non-circular home ranges and the estimation of population density’, Ecology, 100(2), p. e02580.

Egbert, A. and Stokes, A. (1974) ‘The social behaviour of brown bears on an Alaskan salmon stream’, Biology and Management, 3, pp. 40–56.

Elbroch, L. M. and Quigley, H. (2017) ‘Social interactions in a solitary carnivore’, Current Zoology, 63(4), pp. 357–362.

Emmons, L. H. (1987) ‘Comparative feeding ecology of felids in a neotropical rainforest’, Behavioral Ecology and Sociobiology, 20(4), pp. 271–283.

Ganguly, D. and Adhya, T. (2020) ‘How fishing cats *Prionailurus viverrinus* fish: Describing a felid’s strategy to hunt aquatic prey’, bioRxiv, [Preprint].

Gudger, E. W. (1946) ‘Does the jaguar use his tail as a lure in fishing’, Journal of Mammalogy, 27(1), pp. 37–49.

Holt, R. D. (1977) ‘Predation, apparent competition, and the structure of prey communities’, Theoretical Population Biology, 12, pp. 197–229.

Holt, R. D. and Bonsall, M. B. (2017) ‘Apparent competition’, Annual Review of Ecology and Systematics, 48, pp. 447–71.

ICMBIO (2017) ‘Plano de Manejo da Estação Ecológica de Taiamã’, Instituto Chico Mendes de Conservação da Biodiversidade, Brasília, Brasil.

Ivan, J. S., White, G. C. and Shenk, T. M. (2013) ‘Using auxiliary telemetry information to estimate animal density from capture-recapture data’, Ecology, 94(4), pp. 809–816.

Jędrzejewski, W. et al. (2017) ‘Density and population structure of the jaguar (*Panthera onca*) in a protected area of Los Llanos, Venezuela, from 1 year of camera trap monitoring’, Mammal Research, 62, pp. 9–19.

Kantek, D. L. Z. and Onuma, S. S. M. (2013) ‘Jaguar conservation in the region of Taiama Ecological Station, Northern Pantanal, Brazil.’, Publication UEPG. Ciencias Biologicas e da Saude, 19, pp. 69–74.

Kantek, D. Z. et al. (2021) ‘Jaguars from the Brazilian Pantanal: low genetic structure, male-biased dispersal, and implications for long-term conservation’, Biological Conservation, In press.

Long, J. A. and Nelson, T. A. (2013) ‘Measuring dynamic interaction in movement data’, Transactions in GIS, 17(1), pp. 62–77.

Macdonald, D. W. (1983) ‘The ecology of carnivore social behaviour’, Nature, 301, pp. 379–383.

Morato, R. et al. (2018) ‘Resource selection in an apex predator and variation in response to local landscape characteristics’, Biological Conservation, 228, pp. 233–240.

Morato, R. G. et al. (2016) ‘Space use and movement of a neotropical top predator: the endangered jaguar’, Plos One, 11(12), p. e0168176.

Murphy, S. M. et al. (2016) ‘Characterizing recolonization by a reintroduced bear population using genetic spatial capture–recapture’, Journal of Wildlife Management, 80(8), pp. 1390–1407.

Nakano, S. and Murakami, M. (2001) ‘Reciprocal subsidies: Dynamic interdependence between terrestrial and aquatic food webs’, PNAS, 98(1), pp. 166–170.

Polis, G. A., Anderson, W. B. and Holt, R. D. (1997) ‘Toward an integration of landscape and food web ecology: The dynamics of spatially subsidized food webs’, Annual Review of Ecology and Systematics, 28, pp. 289–316.

Rose, M. D. and Polis, G. A. (1998) ‘The distribution and abundance of coyotes: The effects of allochthonous food subsidies from the sea’, Ecology, 79(3), pp. 998–1007.

Royle, J. A. and Young, K. V (2008) ‘A hierarchical model for spatial capture-recapture data’, Ecology, 89(8), pp. 2281–2289.

da Silveira, R. et al. (2010) ‘Depredation by jaguars on caimans and importance of reptiles in the diet of jaguar’, Journal of Herpetology, 44(3), pp. 418–424.

Soisalo, M. K. and Cavalcanti, S. M. C. (2006) ‘Estimating the density of a jaguar population in the Brazilian Pantanal using camera-traps and capture-recapture sampling in combination with GPS radio-telemetry’, Biological Conservation, 129(4), pp. 487–496.

Sunquist, M. and Sunquist, F. (2002) Wild cats of the world. University of Chicago Press.

Tobler, M. W. et al. (2018) ‘Do responsibly managed logging concessions adequately protect jaguars and other large and medium-sized mammals? Two case studies from Guatemala and Peru’, Biological Conservation, 220, pp. 245–253.

Tobler, M. W. and Powell, G. V. N. (2013) ‘Estimating jaguar densities with camera traps: Problems with current designs and recommendations for future studies’, Biological Conservation, 159, pp. 109–118.

Vergara, M (2011) ‘Aspectos ecológicos da onça-pintada (Panthera onca) em uma ilha costeira na região norte do Brasil’, dissertation, Universidade Federal de Santa Maria, Brazil.

